# Leveraging spiking deep neural networks to understand the neural mechanisms underlying selective attention

**DOI:** 10.1101/2020.12.15.422863

**Authors:** Lynn K. A. Sörensen, Davide Zambrano, Heleen A. Slagter, Sander M. Bohté, H. Steven Scholte

## Abstract

Spatial attention enhances sensory processing of goal-relevant information and improves perceptual sensitivity. Yet, the specific neural mechanisms underlying the effects of spatial attention on performance are still contested. Here, we examine different attention mechanisms in spiking deep convolutional neural networks. We directly contrast effects of precision (internal noise suppression) and two different gain modulation mechanisms on performance on a visual search task with complex real-world images. Unlike standard artificial neurons, biological neurons have saturating activation functions, permitting implementation of attentional gain as gain on a neuron’s input or on its outgoing connection. We show that modulating the connection is most effective in selectively enhancing information processing by redistributing spiking activity, and by introducing additional task-relevant information, as shown by representational similarity analyses. Precision only produced minor attentional effects in performance. Our results, which mirror empirical findings, show that it is possible to adjudicate between attention mechanisms using more biologically realistic models and natural stimuli.

## Introduction

Spatial attention is crucial for goal-directed behaviour in many everyday life situations in which one needs to dynamically prioritize processing of information at certain locations in the environment, such as when crossing the street. Spatial attention is generally thought to increase the signal-to-noise ratio of activity in sensory regions representing the attended location. Yet, currently, there are several proposals on how this could be implemented. Some theories propose that spatial attention selectively amplifies the neural signal by changing a neuron’s gain (e.g. Martinez-Trujillo & Treue, 2004; Reynolds & Heeger, 2009), while others posit that spatial attention increases the reliability or precision of processing, thereby emphasizing noise reduction rather than signal amplification as a key mechanism underlying attention’s effects (Feldman & Friston, 2010; Parr & Friston, 2017; Yu & Dayan, 2005). Computational models may allow for arbitration between these different ideas, yet existing models either examined attentional mechanisms in simplified sensory conditions (Feldman & Friston, 2010; Yu & Dayan, 2005) or were not developed to predict changes in performance (e.g. Beuth & Hamker, 2015; Ma, Beck, Latham, & Pouget, 2006; Reynolds & Heeger, 2009; Rothenstein & Tsotsos, 2014). As a result, it remains unclear how spatial attention may facilitate the processing of task-relevant information and thereby performance in more naturalistic settings: through gain modulation, precision modulation (internal noise suppression) or a combination of both.

Deep convolutional neural networks (DCNN) are a way to close this gap in knowledge by linking changes in processing to performance in a fully controlled, yet statistically rich setting (Kietzmann, McClure, & Kriegeskorte, 2019; Richards et al., 2019; Scholte, 2018; Yamins & DiCarlo, 2016). Intriguingly, these networks not only parallel human performance on some object recognition tasks (VanRullen, 2017), but they also feature processing characteristics that bear a lot of resemblance to the visual ventral stream in primates (Eickenberg, Gramfort, Varoquaux, & Thirion, 2017; Güçlü & van Gerven, 2015; Khaligh-Razavi & Kriegeskorte, 2014; Kubilius et al., 2018; Schrimpf et al., 2020; Yamins et al., 2014). Leveraging this link between processing and performance has already enhanced insight into the potential mechanisms underlying shape perception (Kubilius, Bracci, & Op de Beeck, 2016), scene segmentation (Seijdel, Tsakmakidis, de Haan, Bohte, & Scholte, 2020) and the role of recurrence during object recognition (Kar, Kubilius, Schmidt, Issa, & DiCarlo, 2019; Kietzmann, Spoerer, et al., 2019). DCNNs thus provide a promising avenue for systematically investigating how different attention mechanisms may modulate neural processing and thereby, performance.

Here we use a recently developed class of networks, spiking deep convolutional neural networks (sDCNNs, Zambrano, Nusselder, Scholte, & Bohté, 2018), that combine state-of-the-art performance with biologically inspired processing to arbitrate between different proposals of how selective attention may be neurally implemented. Findings from recent studies using DCNNs suggest that changing a neuron’s gain is a viable way to implement selective processing in DCNNs (Lindsay & Miller, 2018; Luo, Roads, & Love, 2020; for a review, see Lindsay, 2020). Yet, these studies did not directly contrast different possible attention mechanisms. Spiking DCNNs as used here provide important additional constraints that can be used to evaluate the feasibility of different mechanisms. Due to their integrated neuron models, information is passed through these networks in temporal spike trains. This makes it possible to measure firing rates, examine population information, and estimate neural latencies and detection times in the network’s output, and then to compare the effects on these measures of different manipulations in relation to findings from neuroscientific studies of attention.

sDCNNs provide two additional advantages compared to DCNNs for studying the mechanisms underlying selective attention. First, due to the neuron models that replace the activation functions used in DCNNs, sDCNN have a more realistic activation regime throughout the network. Commonly, DCNNs use a rectified linear units (ReLU, Nair & Hinton, 2010), while sensory neurons feature an activation that is more sigmoidal-like (e.g. Dayan & Abbott, 2001) and that saturates at high values (Naka & Rushton, 1966). This distinction becomes important when applying multiplicative gain: While for ReLUs there is no difference between the modulation of the in- or output, for sigmoidal activation functions there is a marked asymmetry, which either leads to input or response gain profiles (Ayaz & Chance, 2009; Martínez-Trujillo & Treue, 2002; Reynolds & Heeger, 2009). To understand how selective attention may modulate the gain of neurons, this is a crucial feature that can also help to further situate the findings of earlier DCNN studies using ReLUs for neural processing (Lindsay & Miller, 2018; Luo et al., 2020).

Secondly, sDCNNs feature task-unrelated noise due to their signal transmission properties after spiking conversion. In the brain, neural information transmission also incurs task-unrelated noise for similar reasons (e.g. Allen & Stevens, 1994), which may also affect perceptual performance (Wyart, Nobre, & Summerfield, 2012). One can conceive that a gain modulation might not only boost the signal during processing, but also the noise and can thus have detrimental effects depending on the signal-to-noise ratio. Therefore, having a model that also has task-unrelated noise is crucial for understanding how different attention mechanisms may affect and interact with this signal-to-noise ratio.

In the current study, we capitalized on these properties of sDCNNs to examine how attentional modulation of neural activity may enhance performance. Specifically, we directly compared effects of three kinds of attention mechanisms on performance and network processing, namely: input gain, connection gain and precision. To study the separate effects of input and output gain modulations, exploiting the asymmetry in our activation function, we applied gain to the incoming current of a spiking unit (input gain, Fig. 1A) or we applied gain to the outgoing spike train of the spiking unit. The latter is equivalent to changing the connection strength to the postsynaptic unit (connection gain, Fig. 1B). To model precision, we implemented a mechanism that selectively modulates internal noise, i.e., does not change the gain (precision, Fig. 1C). The effects of these three attention mechanisms were evaluated in the same network during a visual search task using real-world scene images with spatial cueing. Leveraging the full observability of the networks, we also examined the effects of the different attention mechanisms on markers of attentional processing derived from well-established empirical findings in primates, including the magnitude of evoked potentials, firing rates and response latency. In a final step, we systematically examined possible changes in representational content (or the relative change in information present in the network) caused by the three attention mechanisms using representational similarity analysis.

**Figure 1.**
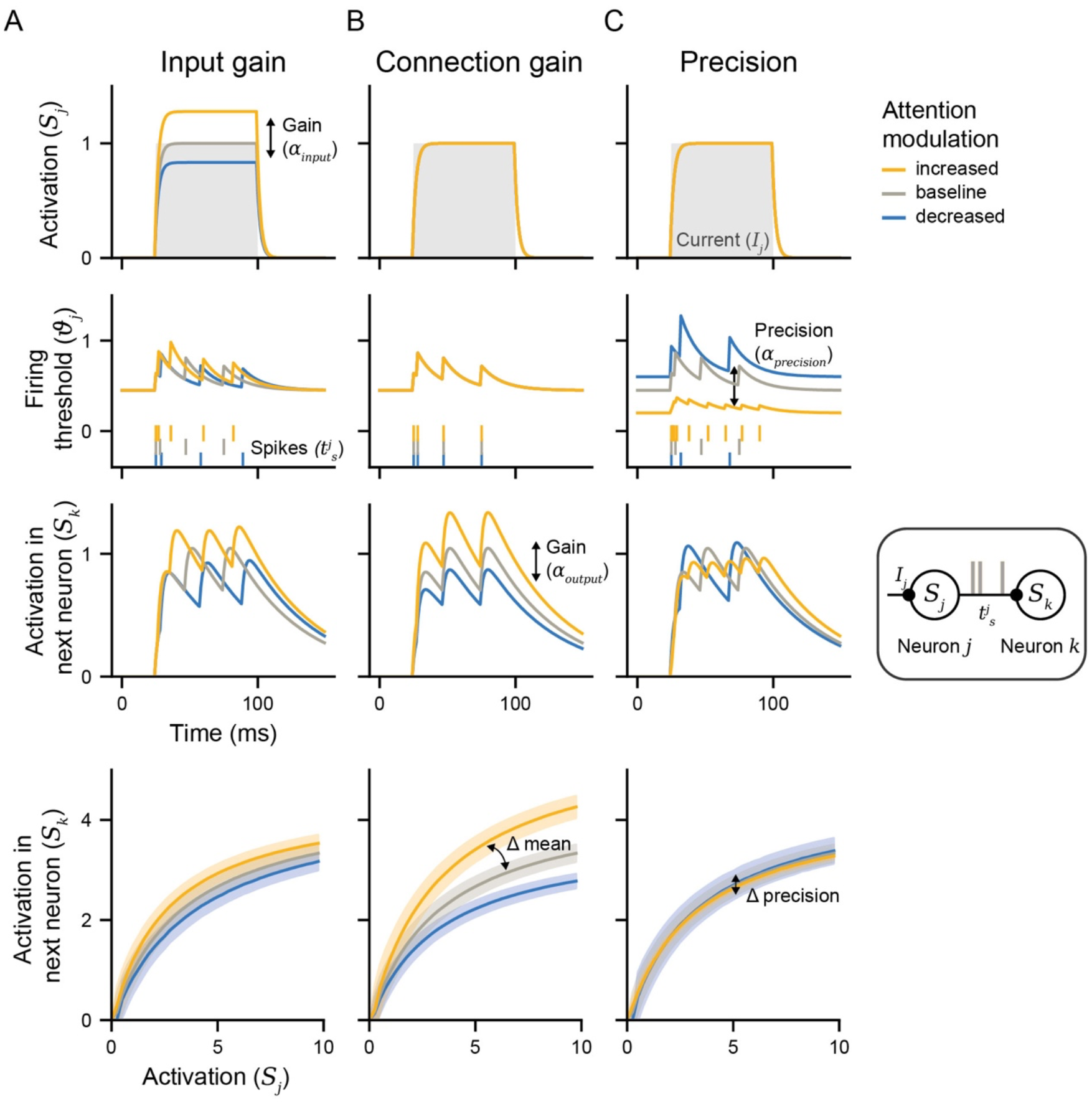
Overview of the three implemented attention mechanisms: input gain, connection gain, and precision. Here, the effects are illustrated in two ways: based on the modulation of the spiking neuron’s processes (top three rows, see inset for a schematic) and based on the activation function, the activity of a spiking neuron over infinite time steps (bottom row). The first three rows show how an incoming current (*I*_*j*_) is converted into post-synaptic activation (*S*_*j*_), which in turn leads to the neuron surpassing its firing threshold (*ϑ*_*j*_) and to produce spikes at times *t*^*j*^_*s*_. This spike sequence is then again transmitted via a weighted connection and integrated by the next neuron, producing the postsynaptic activation *S*_*k*_.(A) For input gain, attentional modulation was implemented by multiplying the integrated current with a spatial weight and a gain factor (*S*_*k*_; top row). This modulation in turn affected all subsequent processes and ultimately led to a modulated firing rate (*ϑ*_*j*_; second row) and a changed activation in the next neuron (*S*_*k*_; third row). Since this manipulation happened before the non-linear process of spike generation, the amplification most strongly affected middle range values in the activation function (bottom row). (B) For connection gain, attentional modulation targeted the postsynaptic weight thereby resulting in an increase of activation in the next neuron (*S*_*k*_; third row) but without changing the spike production (first & second row). In the activation function, this resulted in slope changes, producing the largest modulations for the strongest inputs (bottom row). (C) For precision, attention concurrently modulated the adaptation speed *mf* and the post-synaptic weight. This changed the dynamic firing threshold (*ϑ*_*j*_, second row) and resulted in a change of precision by which the neuron is approximating the input current. By also adjusting the post-synaptic weight, this led to a mechanism that did not affect the mean value but only resulted in differences in the internal noise over time (see modulations of *S*_*k*_, *third row*). The same effect is also illustrated in the width of the shaded areas of the activation function that varies across attention conditions (bottom row).

## Methods & Materials

To directly compare the three different proposed attention mechanisms, we implemented these different mechanisms into the same spiking deep convolutional neural network architecture and assessed their effects on processing and performance during a visual search task with spatial cues.

### Visual search task & dataset

As a first step, we curated a challenging visual search dataset that had a homogenous context, and could contain multiple target objects comparable to naturalistic visual search. In particular, we used street scenes that could contain eight possible target categories as stimuli. Furthermore, we used a dataset with food scenes during model development and for exploratory experiments. To obtain images with a set of potential target categories sharing a context, we first curated these two datasets from the Common Objects in Context database (COCO, Lin et al., 2014). To obtain a homogenous context, we quantified similarity in context as similarity in the stuff-annotations of the COCO dataset (Caesar, Uijlings, & Ferrari, 2016). Specifically, we focused on the super categories from the stuff-annotations (total: 15, e.g., wall, ceiling, sky, water) and defined a vector specifying the presence for all these stuff-annotations for every image. From this, we computed the Euclidean distance between different target category centres (average of all individual vectors belonging to the same target category) so that if a target category on average has the same stuff as another target category, these would have a small distance in such a stuff space. Based on this procedure, we identified a cluster with low distances in stuff-space for street scenes that contained 12 categories (person, bicycle, car, motorcycle, bus, truck, traffic light, fire hydrant, stop sign, parking meter, bench, dog) and for food scenes containing 16 categories (bottle, wine glass, cup, fork, knife, spoon, bowl, banana, apple, sandwich, orange, broccoli, carrot, pizza, donut, cake).

Next, these categories were further processed to select high-quality images with only one recognizable instance of a specific target category present. This is necessary to make the spatial cue informative and advantageous for performance, thus enabling us to quantify spatial cueing in the model’s performance. For the street scenes, we selected images that had a street stuff annotation, for the food scenes, there could be a table, cloth, food-other, vegetable, salad, fruit, and/or napkin stuff annotation. For both datasets, we selected images with target objects that were big enough (*>* 0.05 % of the image), placed in a not too complex scene (spatial coherence *<* 1.2 based on Scholte, Ghebreab, Waldorp, Smeulders, & Lamme, 2009), were not too central (outside of a radius of 5% from the image center) and salient enough (summed object probability density from DeepGaze II *>* 0.04, Kümmerer, Wallis, & Bethge, 2016). This resulted in 8 eligible target categories with at least approximately 50 images with a single target object (street scenes: person, bicycle, car, motorcycle, bus, truck, traffic light, fire hydrant, stop sign, parking meter; food scenes: bottle, wine glass, cup, cutlery, bowl, sandwich, carrot, cake). To obtain a sufficient number of images (around 50 images) for 8 categories in the food dataset, we combined the categories knife, spoon and fork into a single category of cutlery. The code for recreating these datasets is available at https://github.com/lynnsoerensen/SpatialAttention_sDCNN_2020. Due to the datasets’ license status, the final datasets will not be shared publicly but can be requested directly from the authors.

To assess the efficacy of every attention mechanism, each model performed the visual search task on the thus obtained single-target images from the street dataset (*N*^*total*^ = 1628, 224×224 pixels, see Fig. 2C for an example). To quantify the effect of spatial cue validity, we defined a valid spatial cue for every image based on the centre of mass of the target object and an invalid cue pointing to an irrelevant location (Fig. 2B & E). The invalid locations were obtained by randomly sampling from a uniform distribution that was constrained to the minimum and maximum values observed for the valid cues (0.1 - 0.94 for the horizontal and 0.03 - 0.95 for the vertical extent of the image). This sampling process was repeated until the invalid location was at least 0.5 of the image extent away from the valid cue. The valid and invalid locations had on average a distance of 0.62 of the image extent. Due to imperfect COCO annotations, some images also featured two instances of the same target category. To quantify the effect of the spatial bias introduced by the attention mechanisms, we compared performance between a validly cued (centre of mass of a target object in the scene), an invalidly cued (an unrelated location) and an uncued (neutral) processed dataset during most analyses.

**Figure 2.**
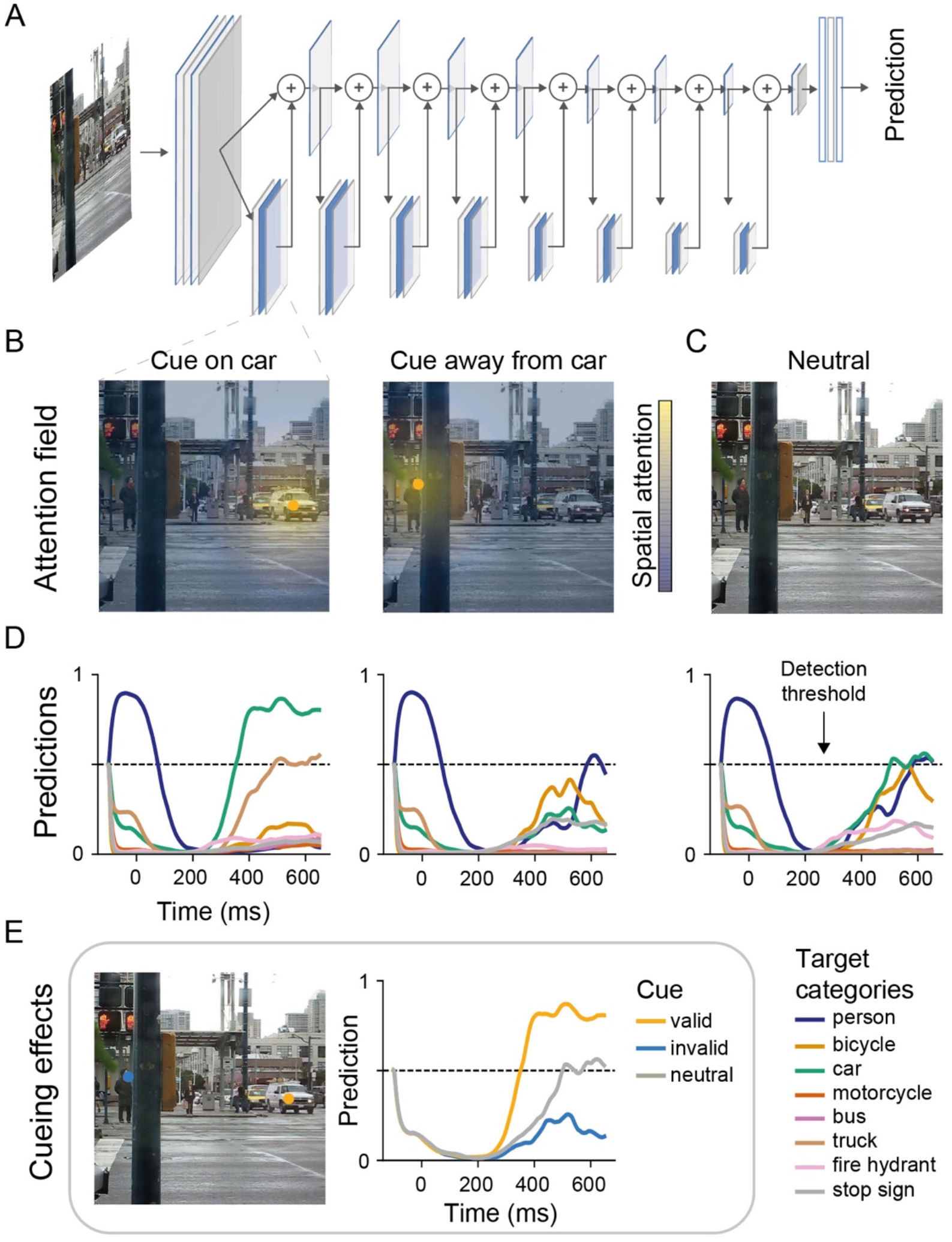
The sDCNN and its naturalistic visual search performance as a function of spatial cueing. (A) An illustration of the ResNet18 architecture with spiking layers depicted by blue frames. The network either takes only an image or also a spatial cue as an input. We implemented the attention mechanisms only in the residual branches of the network (filled blue frames in the lower branch of the network). The output layer was a sigmoid activation function. (B) Illustration of the attention field centering on the cued location, where yellow-coloured regions are allocated more spatial attention and blue coloured less compared to baseline. (C) An example image from the curated dataset of street scenes. (D) Per image and cued location, the sDCNN produces a prediction time course indicating the presence of the eight target categories. An object is present once its prediction exceeds the detection threshold at 0.5. The x-axis shows time relative to the image onset. Passing information through the network takes approximately between 150 - 200 ms due to the spike generation. As a result, the first time points (−100 - 150 ms relative to the stimulus onset) show the biases in the network acquired during training and do not feature any information from the image yet. The left two panels show the network predictions when it was biased by a spatial cue (left: towards the location of the car, middle: away from location of a car). The right most panel shows the neutral predictions for the shown image in (C). (E) Summary of the cueing effects for the car category for the predictions shown in (D). As can be seen, with a valid cue, the network reports the presence of a car more reliably, while it misses the car with an invalid cue. The example image used in B, C and E is licensed under CC BY-NC 2.0 and was obtained from Flickr (https://farm4.staticflickr.com/3326/3259041418_48c260317a_z.jpg).

### Spiking deep convolutional neural networks (sDCNNs)

#### Training & Fine-tuning

We adopted a sDCNN to investigate how different attention mechanisms shape processing and performance. In particular, we used a class of sDCNNs that are comparable to standard DCNNs in most aspects, such as their object recognition performance and training on large-scale database. One of the most important differences is that the standard ReLU activation function is replaced during training with one that captures the activation function of a biologically-inspired neuron model. Specifically, we used a first generation ResNet18-architecture (Fig. 1, He, Zhang, Ren, & Sun, 2015) in which the ReLUs were replaced with such a custom activation function approximating a spiking neuron’s in-and output relationship, which takes the form of a rectified sigmoid-like function. We converted the DCNN to a sDCNN after training and did not directly train a sDCNN because training spiking networks on these deep architectures is exceedingly time-consuming. While it would be desirable to also train a sDCNN in its spiking state, this approach allowed us to yield competitive performance and at the same time still experiment with the properties of spiking neuron models.

We used the network implementation by (Zambrano et al., 2018) and details and derivations of the activation function can be found there. As with standard DCNNs, the network was trained on the ImageNet dataset (Russakovsky et al., 2015) with stochastic gradient descent (initial learning rate: 0.1 with Nesterov momentum of 0.9, decay: 0.0001). The training parameter choices closely followed He al (2015; training augmentation: random cropping and horizontal flipping, test augmentation: center crop). The learning rate was divided by 0.1 every 30 epochs. The final model performed at 64.04% (Top-1 accuracy) on the ImageNet validation set.

For the visual search task, the pre-trained network was fine-tuned on the street dataset by replacing the last fully-connected layer by one with 8 units and a sigmoid activation function. All remaining layers and their weights were kept unchanged. During finetuning, we used images with more than one target object present, selected based on less stringent criteria than the test images (only based on stuff- annontations) resulting in 8640 training and 2160 validation images. The less stringent criteria were necessary to produce a version of the dataset that was large enough for training. The newly added weights were optimized with a binary cross-entropy loss, a learning rate of 0.0001 and an ADAM optimizer for 100 epochs. The final binary accuracy on the multi-object dataset was 88.74%.

#### Spiking inference

After training and fine-tuning, every activation function in the network was replaced with a layer of spiking neuron models that feature membrane potentials, adaptive thresholds and that emit temporal spike sequences as outputs. This means that the trained weights were evaluated with a spiking network that computes continuously over time, thus encoding its activations in binary signals.

The spiking neuron models used here operated on a rate code, and higher firing rates thus encoded higher activation values. This coding principle was implemented by using spiking neuron models that integrate and decay current over time in their adaptive firing thresholds, membrane potentials and refractory responses (Bohte, 2012). Together these features allowed us to convert a continuous signal into a binary signal over time (analog-to-digital converter, (Lazar & Toth, 2003; Yoon, 2017). This step also made it possible to obtain a network performing close to the state-of-the-art (Rueckauer, Lungu, Hu, & Pfeiffer, 2016; Zambrano et al., 2018), while also operating with biologically plausible firing rates. All implementation details of the spiking neurons can be found in (Zambrano et al., 2018) and only the key components are briefly summarised below:

The spiking neuron model consists of four major processes: post-synaptic integration of the incoming spike trains *I*_*j*_ (*t*), their conversion to activation *S*_*j*_(*t*) through a membrane filter, as well an adaptive threshold *ϑ*_*j*_(*t*) and refractory period *Ŝ*_*J*_(*t*), which increase and decay as a function of the timing between emitted spikes.

The post-synaptic current *I* in neuron *j* at time *t* is given by

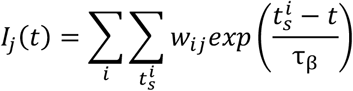

where 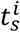 is the timing of the incoming spikes from neuron *i* weighted with *w*_*ij*_. The post-synaptic current decays with the time constant *τ*_*β*_. This becomes the neuron’s activation *S*_*j*_ (*t*) by convolving *I*_*j*_ (*t*) with a normalized exponential membrane filter *ϕ* (*t*):

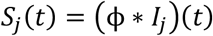

The adaptive threshold *ϑ*_*j*_ is determined by both the resting threshold *ϑ*_0_, the timing of emitted spikes by the neuron, 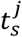, the speed of adaptation *m*_*f*_ and the time constant *τ*_*γ:*_

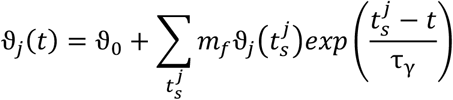

The refractory response *Ŝ*_*J*_(*t*), in turn, is also a function of the adaptive threshold *ϑ*_*j*_ and the timing of emitted spikes 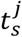 but now decaying with the time constant *τ*_η_:

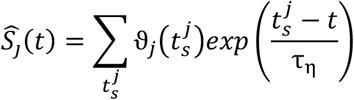

A spike is emitted at time 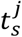 if S(*t*) − *Ŝ*(*t*)> 0.5 *ϑ*_*j*_(*t*), and no spike is produced if that condition is not met, resulting in a binary temporal sequence. This spike sequence is scaled by the constant *h* to correct for the adaptation speed *m*_*f*_ such that the next neuron *k* receives the following post-synaptic current:

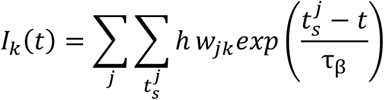

By concurrently adjusting the speed of adaptation *m*_*f*_ and the spike height *h*, it is ensured that the same mean value is approximated by the spiking neuron and thus the trained weights obtained from the non-spiking network are still informative.

In all our experiments, the baseline firing threshold, *ϑ*_0_ and adaptation speed *m*_*f*_ were set to 0.45, *τ*_*γ:*_ was 15 ms and both *τ*_η_ and *τ*_*β*_ were set to 50 ms resulting in a network in which every neuron fired on average at 18.63 Hz. During stimulus presentation, a 100 ms pre-stimulus period was included to take adaptation in the network, such as the saturation of the adaptive threshold, from the image onset response, into account (see Fig. 2D for an example). In total, the network was evaluated over a period of 750 ms. In contrast to the other spiking layers, the sDCNN output layer had a longer membrane potential time constant (*τ*_ϕ_, 50 ms vs. 2.5 ms) and did not produce spikes as output but rather returned its activation *S*_*last*_(*t*), thus producing the smooth prediction time courses as shown in Fig. 2.

All DCNNs, as well as sDCNNs models were implemented, trained and evaluated in Keras with a tensorflow backend. The code is available at https://github.com/lynnsoerensen/SpatialAttention_sDCNN_2020.

#### Attention mechanisms

To compare the different proposed attention mechanisms (input gain, connection gain and precision), we implemented these attention mechanisms into the same base model, thus keeping weights, architecture, and internal noise levels exactly the same.

In line with earlier work (Anton-Erxleben & Carrasco, 2013; Reynolds & Heeger, 2009), we modelled the distribution of spatial attention *R* as a bivariate gaussian distribution over space (Fig. 2D). The centre of the gaussian was placed at the cued location. The standard deviations were kept at 40 pixels for both spatial dimensions. We chose this standard deviation because the average area of an object is equivalent to a circle with a ca. 41-pixel radius. The attention field was normalized to have an average of 0 over all spatial locations based on the assumption that attention involves a redistribution of resources. This thus resulted in some locations being upscaled, while others were downscaled. The spatial reweighting was applied identically for all mechanisms.

The implementation of input gain to a neuron *j* followed

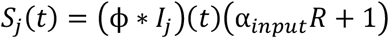

where *R* is the attention field and α_*input*_ is the gain factor. Input gain thus scaled the incoming activation of the neuron and adjusted the spike production accordingly (Fig. 1A).

For connection gain, a gain factor was applied to the outgoing synaptic connection *w*_*jk*_ to the next neuron *k*:

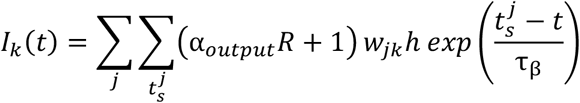

where α_*output*_ is the gain factor. This step scales the out-going spike trains with regard to the impact they will have on the next layer (Fig. 1B). While input gain thus operated on the incoming activation, connection gain targeted the outgoing activation of a neuron by modulating the strength of the connection.

During spiking inference, the binary signals passed between the layers of the network are temporal approximations of the static function learnt during training and these binary signals incur internal noise. Internal noise is thus an unavoidable consequence of using an sDCNN. For implementing precision, we capitalized on this aspect and exclusively changed the internal noise, but not the encoded values, by concurrently adjusting the speed of adaptation *m*_*f*_ and the spike height *h* (Fig. 1C). Specifically, the speed of the adaptation *mf* was modulated by the spatial attention field *R* and the scale factor *α*_*precision*_:

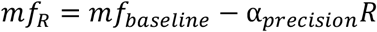

Accordingly, *mf*_*R*_ defines the speed of adaptation for all spatial locations. Such a manipulation also changes the adaptive thresholds in the spiking neuron models, as can be seen in the equations above. When also adjusting the spike height *h* accordingly, this produces a situation, in which the mean approximation of the neuron stays the same, yet the precision of the approximation is varied (Fig. 1C). Depending on the value of *mf*, a neuron thus produced a more or less precise approximation of the output value. As explained above this means that the same underlying function can still be computed, allowing the network to perform, but depending on the spatial attention field *R* and the scale factor *α*_*precision*_ this happens in a more or less precise fashion for different spatial locations. So, in contrast to gain-based mechanisms that change the activation level in the neuron, this mechanism increases the precision, that is, reduces the internal noise, at attended locations, and reduces the precision at unattended locations.

All attention mechanisms targeted the spiking layers in the residual branch of the network (Fig. 2B) and the skip branch was thus unaffected by the attentional modulation. This choice was made based on exploratory experiments on the food dataset (see *Visual search task & dataset*), in which we observed that these branches are set up to be antagonistic, thus modulating the skip branch while also targeting the residual branch cancelled out the effects of attention. In these exploratory experiments, we also observed that it was the most effective to target all residual branches simultaneously. To do this, the attention field was downsampled to match the spatial dimension of the residual branches.

For the main experiments, we compared all mechanisms by quantifying the effect of the spatial bias, that is the difference between a valid, invalid and a neutral cue, introduced by the different attention mechanisms (Fig. 2E for an illustration).

All mechanisms contained a free parameter α. Varying α for all mechanisms effectively reshaped the distribution of attention across space, while keeping the mean identical to the neutral condition (see Fig. 3A for an illustration). For the gain-based mechanisms, attentional modulation varied around a gain factor of 1, and for precision, attentional modulation varied around *mf*_*baseline*_. While for gain-based mechanisms, gain factors above one resulted in enhanced processing, for precision this was achieved with lowered *mf* values relative to *mf*_*baseline*_. Importantly, in contrast to gain-based modulations, precision and the modulation of *mf* was limited by the properties of the spiking neuron model. In particular, adopting very low *mf* values produced too high firing rate regimes, resulting in information loss due to the sampling limit (i.e., the ability to distinguish different spiking sequences, dashed line Fig. 3A).

**Figure 3.**
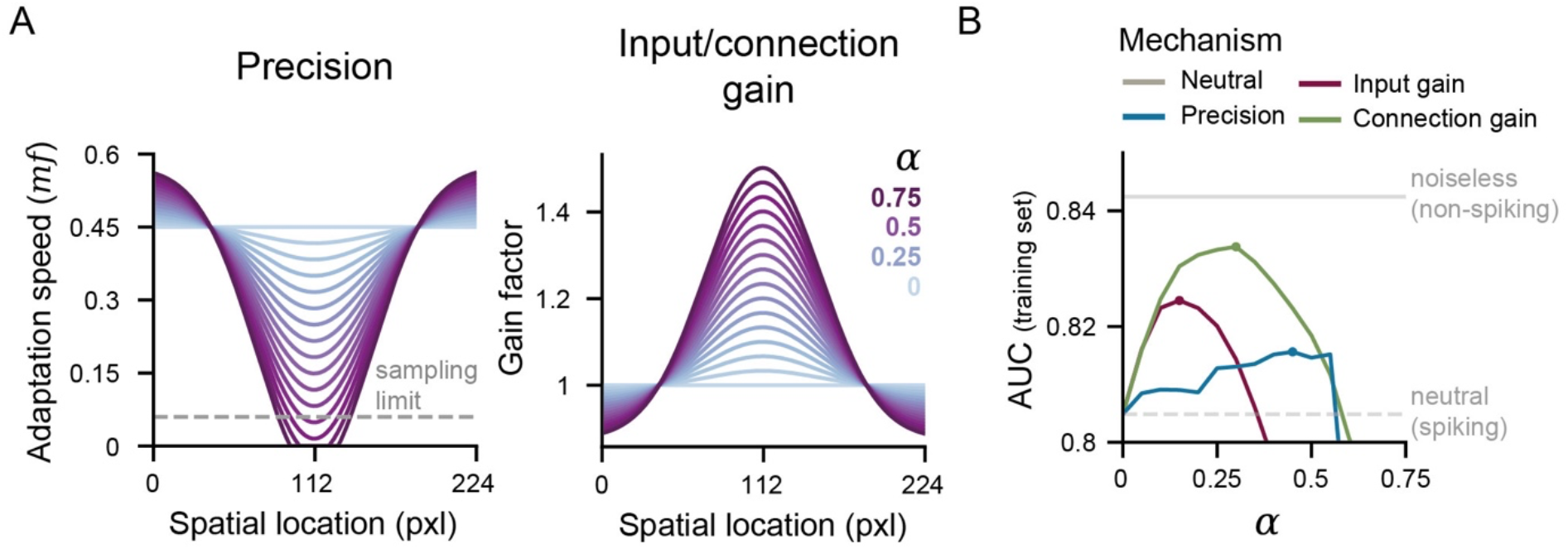
Optimization of attention mechanisms. (A) The impact of α for the three attention mechanisms. Changing α led to the same redistribution of attentional resources across space for all mechanisms. A small value of α approximated the neutral condition (without attentional reweighting), while large values of α led to a strong amplification at the center of the cue and a concurrent suppression in the periphery. For gain-based mechanisms, higher values of α led to higher gain factors at the centre of the attention field. In contrast for precision, increases in α resulted in smaller adaptation speeds at attended locations, leading to increases in firing rates and reduced internal noise. Yet, precision could only be increased up to certain boundary, which was defined by the neuron’s sampling limit. Beyond this limit, the neuron’s signal processing capacity became corrupted. α affected the entire attention field *R*. Here, we only show the central section for illustrative purposes. (B) Identifying the best version for all attention mechanisms. To identify a well-performing version of every mechanism, we performed a grid search by evaluating the benefits in performance derived from a valid cue for all mechanisms on separate training set. This showed that all mechanisms were at their best at different α values, thus benefiting from differently extreme distributions (input gain: 0.15, output gain: 0.3, precision: 0.45, indicated by the same-coloured dots). The sudden collapse in performance of precision beyond α of 0.55 is due to the neuron’s sampling limit.

To identify well-performing versions of every mechanism, we searched through different parameters for α between 0 - 0.75 in increments of 0.05 (α_*input*_, α_*output*_ and α_*precision*_, respectively). Based on the performance on a training set (50% of the single-target dataset, *N* = 814), we chose the best performing model with a valid cue on this data set. While searching the parameter space for α_*precision*_, we observed a rapid decay in performance once *mf* values were reached that surpassed the sampling limit (*mf*_*sampling limit*_ ≈ 0.06, α_*precision*_> 0.6, see Fig. 3). The best performing hyperparameters on the training set were 0.3 for α_*output*_ (connection gain), 0.15 for α_*input*_ (input gain) and 0.45 for α^*precision*^ (precision; see Figure 3B) and these values were adopted for all other simulations.

### Analyses

#### Model performance -target discrimination & detection times

The analyses on model performance were done on the output of the sigmoid activation function obtained from the sDCNN on the held-out single-target images (N=815). We evaluated the model’s target discrimination with the area under curve (AUC) metric across all experiments. To convert the spiking model predictions to this metric, we computed the average of the prediction time-course between 150 and 650 ms after stimulus onset (Fig. 2D). We only evaluated the prediction time-course after 150 ms because the early model responses were largely dominated by the bias terms that the network acquired during training and the saturation of the adaptation in the spiking neuron models (ca. 100 ms before stimulus onset). That responses do not feature much information from the test image yet before this time is due to the temporal characteristics of the spiking neuron models, which require some time to integrate and pass on signals throughout the network hierarchy. As a result, responses to individual images could only be obtained during these later periods.

The attention conditions (valid, invalid, neutral) were statistically compared with permutation-testing for a difference in average AUC (10000 permutations). This was done separately for every mechanism. We performed four pairwise comparisons per mechanism and adjusted our alpha level of 0.05 according to the Benjamini-Hochberg procedure per inference (Benjamini & Hochberg, 1995) for this analysis and for all later analyses. For all analyses, permutation-testing was performed across stimuli, in contrast to model instances, and statistical significance thus implied a significant change caused by a mechanism that is significant given the variability in the dataset.

The model was optimized to predict the presence of a target class by returning values larger than 0.5 due to the use of a sigmoid function at the output layer. We used this feature to estimate the detection time of the model. We defined the detection time as the time point at which the target prediction time course crosses the detection threshold. As described above, prediction time courses were analysed 150 ms after stimulus onset to separate the stimulus response from the general adaptation response caused by the biases in the network (see Fig. 2C for an example).

For both AUC scores and detection times, we estimated 95% confidence intervals using stratified sampling with replacements across stimuli in the test dataset.

Prediction modulation was estimated by subtracting the neutral models’ trials responses (to a single image in the dataset) from the valid or invalid trial responses (to that same image), thereby showing the actual change in prediction introduced by the spatial cue. The 95% confidence intervals were obtained by sampling with replacement from the trials separately per time point.

#### Layer responses – evoked potentials, firing rates & latencies

To determine how observed changes in performance by our attention manipulations may have affected network activity and to relate these to established neural indices of attention, we analysed a layer’s response more carefully. Specifically, we focused on effects on evoked potentials, firing rates and latencies and to this end, analysed the spiking layer of the sixth residual branch (layer 25, 14×14×256). All analyses attempted to follow approaches from neural recording studies as closely as possible.

For all measures, we recorded units representing the centre of mass of the target object for all feature maps in a layer. We chose to do this because we did not want to make a sub-selection of units based on a small sample of stimuli, but instead our goal was to look at the entire layer to get a representative sample. Attention conditions (valid, invalid, neutral cue) were compared by recording from the same set of units.

As outlined above, some attention mechanisms targeted the outgoing synaptic connections *w*_*ik*_, thus measuring the targeted units directly would have the side effect of measuring this very manipulation. We therefore added the spikes of the manipulated unit into another unweighted unit.

For the evoked potentials, we integrated the spikes from the manipulated neurons and read out the potential. These are thus a direct precursor of the firing rates. For the firing rates, we reported the output of the unweighted neuron. For illustration purposes, the obtained spike histograms across all features maps were smoothed with a temporal gaussian kernel (standard deviation of 8 ms) in Fig. 5B.

The response latency for the spike responses were calculated by a metric that closely followed (Lee, Williford, & Maunsell, 2007; Sundberg, Mitchell, Gawne, & Reynolds, 2012). Specifically, latency was defined as the time by which the spike density function reached 50% of the maximum firing rate of the first peak in the transient response after stimulus onset.

To estimate the latency, the spike responses were first smoothed over time with a gaussian filter of 8 ms. In a next step, every trial was compared back to baseline measurements in which 16 average images based on 50 randomly picked images from the training set were presented to the network. Based on the activation of the baseline activation in every trial, we determined the 99.99% percentile of activation (corresponding to 3.72 SEM in Sundberg et al., 2012) and used this value as a criterion to identify the first local peak after stimulus onset. Both baseline and experimental trials were baseline-corrected based on the 50 ms prior to image onset.

If a trial did not surpass the criterion for activation, we could not estimate the latency, which led to a different number of trials that were excluded per attention mechanism and cue (see Table 1) of a total of 815 trials for every cue condition. For all measures, we estimated 95% confidence intervals by resampling with replacements across trials.

**Table 1.** Missing trials for latency estimation per attention mechanism and cue. The procedure of the latency estimation involved to assess whether a spiking unit was more active compared to baseline. If this baseline criterion was not exceeded, no latency estimate was made. More active networks are thus less likely to have missing trials compared to inactive networks.

| **Mechanism** | **Valid cue** | **Invalid cue** |
| --- | --- | --- |
| Precision | 55 | 112 |
| Input gain | 34 | 96 |
| Connection gain | 25 | 96 |
| Baseline (no cue) | 133 |  |

#### Representational similarity analysis

We next looked at how our different attention mechanisms and conditions may have affected the information in our network using representational similarity analysis. Due to computational constraints, we used a subset of the images compared to the rest of the experiments (400 in total, 50 per category). The resulting sub-dataset was classified by the noiseless model with a similar level of performance as other randomly drawn samples from the dataset (AUC: 0.8452 vs. 0.8409, selected vs. resampled).

For every image, mechanism and attention condition, we obtained the evoked activations for network layer 37. We chose this layer because it is the very last spiking layer of the ResNet blocks and is not targeted by attention itself since it is part of the skip branch.

We obtained the pairwise Pearson correlation between every image for every time point and model with the respective mechanism and attention condition separately, thus giving us seven representational dissimilarity time courses.

The goal for this analysis was to understand how the representations in the network are altered by the different attention mechanisms. We reasoned that a closer similarity to the non-spiking, noiseless network means that the spiking network recovers from the introduced noise due to the spatial attention cue. In a similar vein, we expected that the similarity with the categorical RDM should grow if there is additional target object information present that helps the network at the task. To test this, we correlated every timepoint of the seven RDM time courses with either the noiseless or categorical RDMs.

The non-spiking, noiseless network’s RDM and the categorical RDM are correlated because the noiseless network has been optimized to distinguish these object classes and the categorical RDM embodies the best possible distinction between the object classes, the ideal observer. Yet, these RDMs are also not identical. For example, the noiseless RDM might contain systematic errors made by the network due its imperfect solution of the task found during training. Observing that this kind of variation (i.e., systematic errors acquired during training) increases due to selective attention is a unique signature of the noiseless network and indicative of an attention mechanism that reduces the noise on the distinction learned during training, a scenario we term noise suppression. Conversely, if the RDM affected by an attention mechanism contains more categorical information than the categorical information of the noiseless network, this indicates that the selective attention mechanism increased the distinction between the object categories beyond the trained weights. This would suggest that new categorical information was added and thus that the signal in the network was enhanced. In our analysis, importantly, we could dissociate between these two scenarios. Specifically, to this end, we calculated the partial correlation for these two predictors, thus giving us the unique contribution of both predictor RDMs, while keeping the influence of the other predictor constant. To compare this to the neutral model, we subtracted the partial correlations of neutral condition from all other partial correlations. We assessed statistical significance by comparing the mean bootstrapped difference between 150 - 650 ms to zero. For all time courses, we estimated 95% confidence intervals by resampling with replacements across the RDMS per timepoint.

#### Software

In addition to custom code (https://github.com/lynnsoerensen/SpatialAttention_sDCNN_2020), the results presented here were obtained while relying on the following Python packages: NumPy (Harris et al., 2020), keras (Chollet, 2015), TensorFlow (Abadi et al., 2016), Pandas (McKinney & Others, 2010), Pingouin (Vallat, 2018), Scikit-Learn (Pedregosa, 2011) and SciPy (Virtanen et al., 2020). Data visualization was done using matplotlib (Hunter, 2007) and, in particular, seaborn (Waskom et al., 2020).

## Results

We adopted a sDCNN to investigate how different attention mechanisms affect processing and performance during a challenging visual search task.

### Connection gain most effectively produces spatial cueing effects on performance

Human subjects are faster and more accurate when targets occur at a validly cued spatial location compared to an invalidly cued one (Carrasco, 2011; Posner, 1980). In our first analysis, we show that connection gain is best at changing the network’s performance in the same way.

Fig. 4A shows the prediction modulation for all attention mechanisms (valid/invalid vs. neutral trial predictions) evolving over time. We see that for all mechanisms, the target class is modulated up or down depending on the cue’s validity, suggesting that the spatial cue led to a modulation of the target class predictions and thus came at a cost or benefit to the model’s prediction. To quantify the effect on target discrimination of this modulation per mechanism, we computed the area under the curve scores (AUC) and compared these scores to those of a neutral (without any spatial attention bias) and noiseless, non-spiking network (as obtained after training and before spiking conversion). For all networks, we inspected the average prediction in the period of 150 - 650 ms after stimulus onset. To assess whether the mechanisms introduced spatial cueing effects, we performed pairwise permutation tests contrasting target discrimination in validly and invalidly cued trials. We found that only connection gain produced different levels of target discrimination as a function of cue validity (*p* = .002) that were larger than the variability expected from the stimulus set (Fig. 4C).

**Figure 4.**
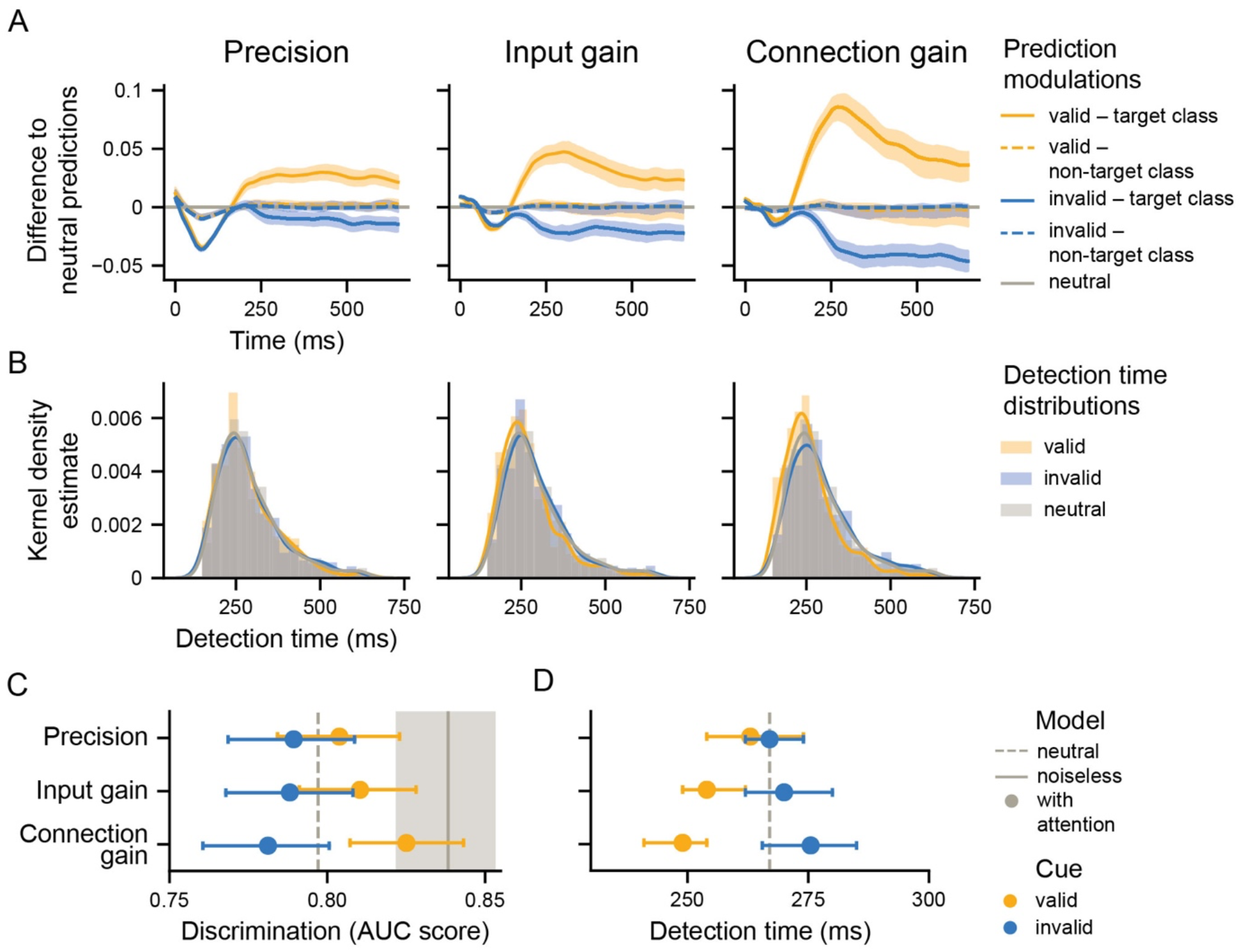
Connection gain most strongly modulated performance. (A) Illustration of how the network’s average responses were modulated by the spatial cues by subtracting the attended trials from the neutral trials. The panels show the data for each attention mechanism. (B) Detection time distributions across the different trial types for all mechanisms. (C) Overview of spatial cueing effects on target discrimination (x-axis) for the different attention mechanisms (y-axis). The y-axis is ordered according to valid cue performance. The shaded area around the noiseless model as well as the error bars represent the 95% confidence interval resampled across dataset stimuli. The attention mechanisms were defined based on a prior grid search over the gain parameter on a separate data set (see Fig. 3). (D) Overview of spatial cueing effects on detection times for all attention mechanisms.

**Figure 5.**
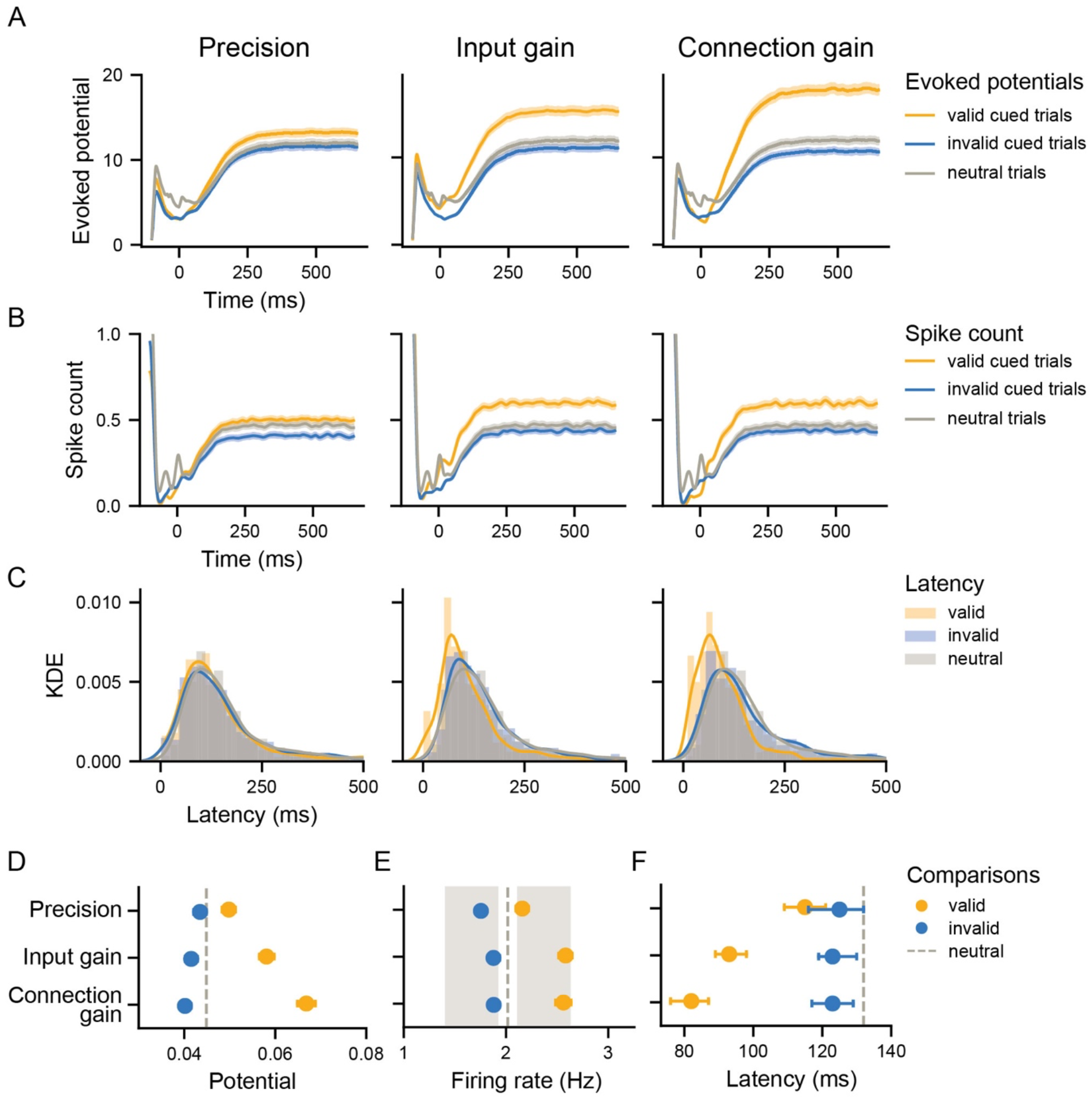
Effects of precision, input and connection gain on key neural indices of attention: evoked potentials, firing rate and latency. (A) Evoked potential time courses measured from spiking units at the centre of the target object in a given image for the three different conditions (valid, invalid, neutral cue) for all mechanisms. Shaded areas depict the 95% confidence interval (CI) obtained with resampling across stimuli. (B) Average spike count time courses from the same units shown in (A). The spike count time courses have been temporally smoothed with a gaussian kernel. (C) Latency estimate distributions for all mechanisms. Latencies were estimates from the smoothed spike count histogram and defined as the time when 50% of the first peak was reached. KDE stands for kernel density estimate. (D) Mean evoked potential between 150 - 650 ms compared across different attention mechanisms. The error bars depict the 95% CI for D, E and F. (E) Mean firing rates for the different attention mechanisms. The grey boxes indicate a modulation of 5 - 30% from baseline as observed in experimental data (Maunsell, 2015). (F) Median latencies for the different conditions across attention mechanisms.

Selective attention normally (in the brain) operates in the context of noise. It is hence not surprising that in the sDCNNs, that feature task-unrelated noise, target discrimination was reduced compared to the noiseless network (Fig. 4C). To assess how the different attention mechanisms can deal with noise, we tested for a difference between all mechanisms with a valid cue and the noiseless model’s target discrimination. We found that only for precision target discrimination was significantly reduced compared to the noiseless network (*p* = .01). This thus indicates that especially gain-based mechanisms were able to overcome the noise inherent in sDCNNs with a valid cue.

As a next step, we evaluated if spatial cueing also influenced detection time. The sDCNN were trained to report the presence of a target category by using a sigmoid function with a cut-off at 0.5 during a multi-label task. We here interpret the first time point after 150 ms at which this threshold is passed as detection time. Fig. 4B shows the detection time distributions for all studied mechanisms for the different trial types. Comparing the medians of these distributions between the valid and invalid cues reveals significant differences in detection times for both gain-based mechanisms (all *p <* .002), but not for precision (*p* = .359, Fig. 4D). Again, a model with connection gain produced the largest reaction time validity effects.

Taken together, both the model’s ability to discriminate as well as its detection times were most strongly modulated by connection gain, reproducing the characteristic shifts in performance observed in spatial cueing studies in humans.

### All attention mechanisms can qualitatively replicate neural changes

The effects of spatial attention on processing in visual cortices have been extensively documented (Maunsell, 2015). Here, we investigated how the different mechanisms implemented in our sDCNNs induced neural changes of the kind and magnitude observed in empirical studies. In particular, we examined changes in evoked potential, firing rates and latency. For this we looked at spiking units from the sixth ResNet block, which is in the middle of the network (Fig. 2A). These units still have some degree of spatial specificity while also having a sizable impact on network performance. Within the sixth block, we selected units that represented the features at the centre of mass of the target object. We recorded from these units under three conditions: without a cue, with a valid cue, and with an invalid cue.

The evoked potentials, that is, the integrated currents inside of the spiking units, are the most comparable to the activations observed in non-spiking DCNNs. If an input gain mechanism works as expected, it should modulate these values, as these are the proportional markers of encoded activation. With the sDCNN, we have the possibility to see how the encoded values at valid and invalid locations change with the different attention mechanisms, and to link these changes directly to changes in firing rate. In contrast, measuring the evoked potentials in single neurons simultaneously in a large population can be experimentally challenging. To obtain the evoked potential without measuring the direct consequence of the attentional manipulations (e.g., a change in post-synaptic weight) as well, we integrated the spike trains into another neuron.

Fig. 5A shows the average evoked response across the entire dataset (*N* = 815) for the three attention mechanisms. Comparing the mean modulation between 150 - 650 ms across mechanisms revealed that the evoked potentials were modulated significantly by all attention mechanisms, as indicated by significant differences in potential amplitude between the valid and invalid cue conditions (all *p <* .001,

Fig. 5D). Following our behavioural results, connection gain numerically had the greatest effects on evoked responses, and precision the smallest.

Next, we studied how these changes in evoked potential may translate to changes in firing rate. Modulation of firing rate by spatial attention is a very common observation in electrophysiological studies. Firing rates typically change as a function of cue validity with a modulation range of 5 - 30% compared to baseline (Maunsell, 2015). Here, we found that all mechanisms resulted in changes within this range.

For measuring the firing rates, we recorded spike trains at the output of the next unit. Since for connection gain, the strength of the connection to the next neuron was manipulated, this kept the impact of attention for all measurements the same. Fig. 5B shows the average response across the dataset for all mechanisms. In contrast to typical analysis protocols of experimental data, we did not make any preselection of units based on their responsiveness. Because our networks are sparse in their activations, this results in low firing rates (Fig. 5B).

Across mechanisms, there was a significant increase in firing rates in response to a valid compared to an invalid cue (all *p <* .001, Fig. 5E). To link these changes back to experimental data, we plotted the range of 5 - 30% of modulation compared to the neutral model (grey shaded areas). It becomes clear that all models are producing changes that are within the biologically observed range with connection gain being the best performing mechanism. Interestingly, the two gain-based mechanisms were associated with comparable firing rate modulations.

Lastly, spatial attention has not only been shown to alter the response magnitude of visual cortical neurons, but also to modulate the latency of their responses. (Sundberg et al., 2012) reported that attention was associated with a reduction in latency between 0.5 - 2 ms of both the spiking and LFP responses of neurons in V4. Similar findings have also been reported for MT, suggesting an overall reduction in response latency across multiple visual areas (Galashan, Saßen, Kreiter, & Wegener, 2013). In our next analysis, we examined the latency changes introduced by the different attention mechanisms in response to a valid and invalid cue. In brief, we find that all mechanisms markedly affected the processing latency, yet at a much larger magnitude than observed in neural data and that this change was mainly driven by benefits in response to a valid cue.

To obtain latency estimates, we re-analysed the firing rate data with regard to the first modulation compared to baseline activity per trial. We defined latency as the time point by which the smoothed spike density function reached 50% of the maximum firing rate of the first peak in the response after stimulus onset following Sundberg et al. (2012).

Fig. 5C shows the distribution of estimated latencies for all mechanisms. The difference in distributions along the x-axis indicates a large decrease in latency for valid cues. Indeed, for all attention mechanisms, we observe faster response latencies in valid compared to invalid cue conditions (all *p <* .033, Fig. 5F).

Yet, only valid cues produced a reliable latency benefit across all mechanisms compared to the neutral condition (all *p <* 0.001). Again, following our behavioural results, latency reductions were largest for connection gain, and smallest for precision. Furthermore, observed latency differences were much larger than those typically observed in neural processing (11.5 - 44 ms vs. 0.5 - 2 ms). This suggests that there are potentially important differences between the neural and sDCNN firing rate data in the onset response and the temporal adaptation that may have made this analysis approach less suitable for sDCNNs.

In sum, we found that all mechanisms modulated evoked potentials, firing rates and latencies according to cue validity, thereby mostly paralleling observations from neural recordings.

### Only gain mechanisms introduce additional information

Lastly, we sought to capture the impact that these various attention mechanisms have on the representations in the network. In primates, spatial and category-based attention have been reported to improve neural population coding in inferior temporal cortex of real-world objects among distractors (McKee, Riesenhuber, Miller, & Freedman, 2014; Zhang et al., 2011), suggesting that selective attention can amplify information in a noisy neuronal population. What still remains an open question is whether this improvement in population coding stems from a less noisy representation (internal noise suppression) or rather can be attributed to additional information about the attended object or location (signal enhancement). Therefore next, we aimed to understand how specifically the different attention mechanisms increased the SNR. Using a representational similarity analysis, as detailed below, we show that gain-based mechanisms achieved this by adding signal to the network’s representations, while precision resulted only in noise suppression.

With our models, we are in the position of having full access to all the units in a layer without incurring measurement noise. In addition, we also have a notion of the noise of the network at baseline (neutral network) and the representation of the noiseless network (non-spiking). With this, we can disentangle the relative contribution of noise recovery and added signal due to the spatial information of the attention cue by using representational similarity analysis (Kriegeskorte, Mur, & Bandettini, 2008). For this analysis, we define noise recovery as an increase in similarity between a spiking network and its noiseless counterpart (Fig. 6B), and an increase in signal as an increase in similarity of the spiking network to a fully categorical representation (Fig. 6A).

**Figure 6.**
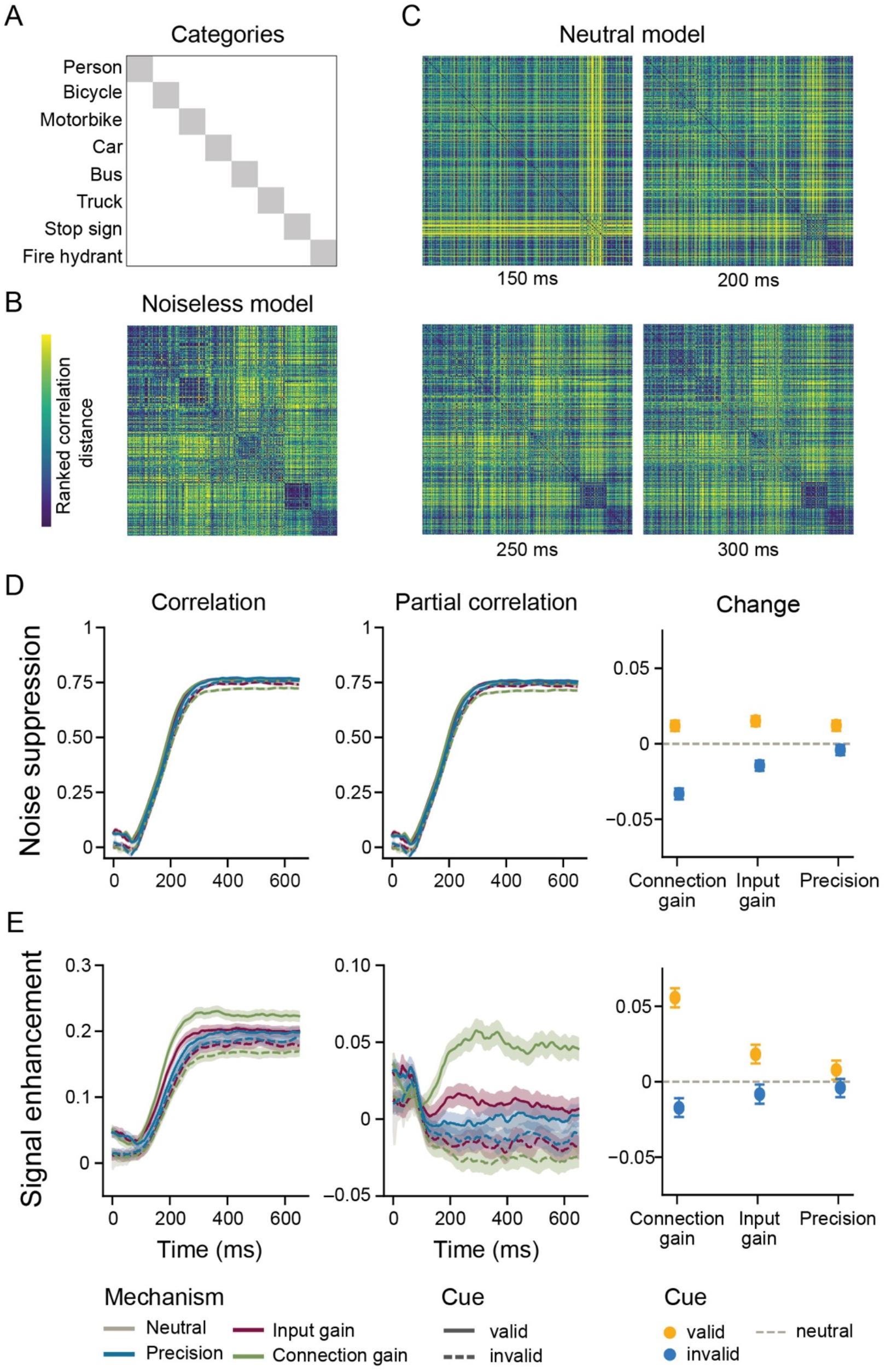
Gain mechanisms selectively enhance categorical information. (A) Organization of the categories in the representational dissimilarity matrix (RDM). For every category, there were 50 images. The categories were organized according to visual similarity. For illustration purposes, the correlation coefficients were ranked. (B) RDM of the layer of interest from the noiseless, non-spiking network. (C) RDMs from four timepoints after stimulus onset for the neutral network. (D) Left panel: Pearson-correlation coefficients between the noiseless model and the temporal RDMs separately for the three attention models (precision, input gain, connection gain) and the neutral model. The shaded areas represent the 95% confidence interval obtained from resampling across stimuli. Middle panel: Partial correlation coefficients for the comparison in the left panel and now controlling for the categorical RDM shown in (A). Right panel: Average difference in partial correlation between attention models and the neutral model. (E) Left panel: Pearson-correlation between the categorical model and the temporal RDMs shown separately for the three attention models and the neutral model. Middle and right panel: Same as in D, but now controlling for the noiseless model.

Due to computational constraints for the sDCNN, we selected a random set of 50 images per category (400 images in total). Since the effects on population coding have been reported in inferior temporal cortex, we chose the first spiking layer of the decoding block (Fig. 2A, the first spiking layer after the last ResNet block). We constructed a representational dissimilarity matrix (RDM) for every time step separately for each network (attention mechanism) presented with a valid or invalid cue. Fig. 6C shows four example time frames from the neutral network to illustrate the temporal evolution of the RDMs. To quantify noise suppression and signal enhancement, respectively, we used the correlation with the noiseless (Fig. 6B) and categorical model (Fig. 6A) as our metric. Since both of the RDMs are highly correlated, we disentangled their unique contributions by computing the partial correlation. We used this metric to estimate whether there was a statistically significant shift in explained variance by comparing the unique correlations for a valid and invalid condition to the neutral condition, respectively. Within a given mechanism, we constructed a 95% confidence interval for the difference between the valid and invalid trials, and estimated variability by means of bootstrapping with replacement across stimuli.

Examining the changes in RDMs over time across the different mechanisms and cues, we see an increasing correlation to the noiseless network over time that stabilizes around 300 ms after stimulus onset (Fig. 6D, left panel). Comparing the three different attention mechanisms to the neutral network after controlling for categorical information (Fig. 6D, middle panel), we observe that all mechanisms became more similar to the noiseless network upon presentation with a valid cue (all *p <* .002). When examining the effect of an invalid cue (Fig. 6D, right panel), only input gain and connection gain led to a decrease in similarity (all *p <* .002, *p*_*precision*_ = .02). Pairwise comparisons within a validity condition indicate that all mechanisms were different from one another (all *p <*.002) except for precision and connection gain with a valid cue (*p* =.836). Altogether, this suggests a reinstatement of the noiseless network representations for all mechanisms if presented with a valid cue, and a decrease in similarity for invalid cues for gain mechanisms.

For signal enhancement as expressed in a correlation with the categorical model, we observed similar temporal characteristics as for the noise recovery (Fig. 6E, left panel). Yet, after controlling for the effect of noise recovery, we found that only the mechanisms that changed the signal (i.e., input and connection gain) featured significantly more additional categorical information with a valid cue (Fig. 6E, middle & right panel, *p*_*precision*_ = .012, remaining *p <* .002). The same was true for an invalid cue, where only input gain and connection gain, but not precision, decreased the amount of categorical information compared to the neutral network (*p*_*precision*_ = .012, remaining *p* < .002). Pairwise comparisons between the mechanisms within a validity condition further support the existence of differences in signal enhancement, as significant differences between all mechanisms were observed for both valid and invalid cues (all *p’s* < .002).

In sum, the different attention mechanisms introduced different representational changes: While all examined mechanisms reinstated the representations used by the noiseless network to a similar degree, gain mechanisms that targeted the signal were more effective at adding categorical information beyond the trained weights compared to the noiseless network. This suggests that additional categorical information was derived from the spatial cue that effectively helped performance.

## Discussion

In this modelling study, we examined how spatial attention may affect sensory processing and performance on a challenging visual search task using a sDCNN. Specifically, we directly contrasted effects of three different mechanisms previously proposed to subserve selective attention (e.g. Dayan, Kakade, & Montague, 2000; Feldman & Friston, 2010; Martinez-Trujillo & Treue, 2004; Reynolds & Heeger, 2009), namely gain modulation on the input to a neuron, gain modulation on its post-synaptic connection and modulation of the neuron’s precision (internal noise). We found that connection gain was most effective at implementing spatial attention, as indicated by the largest performance modulations, whereas precision and input gain were less effective. That is, connection gain modulations produced the largest difference in detecting and discriminating targets occurring at a validly cued spatial location compared to an invalidly cued one, a pattern commonly observed in humans (Carrasco, 2011; Posner, 1980). Moreover, connection gain also well reproduced several key experimental findings in visual cortex (Maunsell, 2015), including a proportionate modulation in firing rates. Disentangling the representational changes introduced by the three main mechanisms in the network revealed that gain-based mechanisms in particular added task-relevant information, while all networks showed similar recovery from the noise. These results mirror findings from the animal and human literature that show that attention can enhance the representational content of neural activity (e.g. Jehee, Brady, & Tong, 2011; Zhang et al., 2011). Together, these findings advance our understanding of how spatial attention might be mechanistically implemented at the neural level, as discussed in more detail below.

Our finding that connection gain was not only more effective than precision, but also than input gain, highlights that the asymmetry in the activation function plays a big role in the efficacy of gain on the model’s performance: connection gain was more effective at enhancing activation in a useful fashion and thereby biased the network’s performance more strongly compared to input gain. This result helps us to better understand how this asymmetry can act as a constraint for a gain mechanism. This finding also sheds new light on past studies that used ReLUs to implement gain (Lindsay & Miller, 2018; Luo et al., 2020): A shared feature between their gain implementation and our connection gain is that both approaches specifically boost larger values, that is, higher values are proportionally more affected compared to smaller values. Based on our results, we can thus speculate that these larger values in particular are important for boosting performance, at least in DCNNs.

In contrast to gain, we found that precision was not as effective as gain-based mechanisms at implementing attentional selection as reflected in performance measures. This finding was unexpected since global changes in precision have been shown to lead to substantial changes in noise during processing in a previous sDCNN study and are a powerful way to increase and decrease the network’s performance (Zambrano et al., 2018). It also does not align with notions that assign an important role to reliability or precision in selective information processing, and that emphasize noise reduction rather than signal amplification (Dayan et al., 2000; Feldman & Friston, 2010; Parr & Friston, 2017). One potential explanation that could reconcile our finding with this past work is that we did not use precision in the context of a Bayesian observer framework (Feldman & Friston, 2010; Parr & Friston, 2017). Instead, our notion of precision is directly grounded in the signal-detection framework, where a signal is represented in a more or less narrow distribution, making it thereby more or less distinguishable from noise. It is a possibility that such a mechanism proves to be more effective when used on prediction errors rather than on stimulus-driven information. Precision may also play a more dominant role in well trained systems: a set of event-related potential (ERP) studies by (Itthipuripat, Cha, Byers, & Serences, 2017; Itthipuripat, Ester, Deering, & Serences, 2014) showed that spatial attention was associated with a gain modulation of early visual-evoked potentials early in training, but with noise reduction at advanced stages of training (Itthipuripat et al., 2017). These findings suggest that the mechanisms through which spatial attention facilitates performance may depend on the specific behavioural training regime used, with gain-type mechanisms subserving selective information processing in relatively untrained systems, as in our models. Under the assumption that learning affects the synaptic strength akin to a momentary gain change, we can make the prediction that precision should be especially successful in our models once we allow for weight adaptation after trials with attentional selection. Future studies are necessary to test this prediction.

To accommodate our studied attention mechanisms, we augmented a standard DCNN with spiking neurons, resulting in three key changes: a changed activation regime due to the sigmoidal transfer function, temporal processing, and internal noise during spiking inference. While the first change was essential to define input and connection gain and to address the issue of saturation in biological neurons, the latter enabled us to implement precision and to mimic noisy neural transmission. The second change, temporal processing, was useful to connect our model to both behavioural reaction times and neural latencies. Interestingly, temporal processing in our model, which was solely obtained by combining the neuron model with standard feedforward network weights, already exhibited dynamics attesting to evidence accumulation over time (see Fig. 2D). This highlights how our approach can enrich the standard DCNN performance measures, for instance, as a baseline model for temporal dynamics and speed-accuracy trade-offs for other more complex temporal vision models deploying recurrency (e.g. Spoerer, Kietzmann, Mehrer, Charest, & Kriegeskorte, 2020). For the activation regime, our results establish that the asymmetry between input and connection gain can be a decisive factor for the efficacy of an attention mechanism and that competitive task performance can be maintained despite this design choice (Zambrano et al., 2018). Indeed, some of our framework could be pursued without the spiking neurons and one could adopt a feedforward DCNN with an altered activation function instead as we have used during training. While sacrificing a level of detail, such a network would still permit to further explore the efficacy of attentional gain modulations in a leaner computational setting.

The current results are based on a ResNet-18 architecture, and we chose this architecture because it has been shown to be successful at object recognition while having a relatively modest number of parameters, a desirable property when implementing memory-intensive spiking networks. The use of this particular architecture may have affected our results. For instance, it is conceivable that the interplay between the residual and skip connections typical for ResNets (see Fig. 2A for an illustration) was particularly suitable for gain-based mechanisms in contrast to precision. While we a priori did not anticipate such an interaction between architecture and attention mechanisms, follow-up studies with simpler feedforward architectures are necessary to further examine the extent to which our findings generalize to other network architectures.

Our findings on input and connection gain raise the question of how these mechanisms might link to selective attention in biological neurons. The effects of input gain could be related to what has been referred to as contrast gain in neural processing (Reynolds, Pasternak, & Desimone, 2000). Such neural changes are mathematically equivalent to a multiplication of the incoming current (Reynolds & Heeger, 2009). Yet, how specifically this multiplicative gain could be implemented in a neuron is still matter of active research (for a recent review, see Ferguson & Cardin, 2020). The effects of connection gain resemble neural changes described as response gain: increased firing rates that scale beyond the maximum response observed under neutral conditions (McAdams & Maunsell, 1999; Treue & Martínez Trujillo, 1999). This gain profile may arise from the same neural populations as contrast gain, yet in situations in which a relatively small attention field is paired with a large stimulus (Reynolds & Heeger, 2009). For its biological implementation, it has been suggested that both neural contrast and response gain can be obtained by combining a multiplicative gain on excitatory inputs with lateral or feedforward inhibition (Beuth & Hamker, 2015). Another recent proposal is the addition of a current in a recurrently connected excitatory-inhibitory circuit, which results in a multiplicative gain that can also show a switch from contrast to response gain (Lindsay, Rubin, & Miller, 2019). An alternative for how connection gain effects could be implemented independently of contrast gain is as a change in synaptic efficacy, as was reported in a set of studies in LGN and V1 (Briggs, Mangun, & Usrey, 2013; Hembrook-Short, Mock, Usrey, & Briggs, 2018). Future experimental as well as modelling studies will be crucial to further link biological circuits to computational functions during attentional selection. Our results suggest that there is a computational advantage in amplifying a neuron’s outputs, akin to both response and connection gain, rather than its inputs, even when the system features noise.

A central goal of the current modelling experiments was to study attention mechanisms in the context of more biologically plausible processing constraints. A challenge inherent to any such endeavour is that the validity of findings is determined by the accuracy of the model in capturing relevant biological properties. Our study was conducted with adaptive spiking neurons (Bellec, Salaj, Subramoney, Legenstein, & Maass, 2018; Bohte, 2012). We chose this neuron model because it allowed us to implement key biological properties (e.g. adaptive thresholds, saturation at realistic firing rates, (Zambrano et al., 2018) and relate changes in firing rates, population information, neural latencies, discrimination performance and detection times in the network’s output to key findings from behavioural and neuroscientific studies of attention. While thereby providing important additional, neurally grounded constraints for evaluating the feasibility of different mechanisms compared to existing DCNNs that have used ReLUs, our neuron models do not have stochastic firing thresholds or other sources of inter-trial variability due to the intended conversion after training. As a consequence, it is not possible to relate our results to effects of selective attention on spike count variability or noise, that have also been reported in the literature as a measure of noise (Cohen & Maunsell, 2009; Mitchell, Sundberg, & Reynolds, 2007, 2009). A recent study suggests that such measures can be reproduced in a circuit of neuron models injected with synaptic noise (Lindsay et al., 2019). Future studies that include such synaptic noise and also assess the noise covariance between neurons are necessary to establish a more direct link to noise correlations and to complement the here presented results.

Finally, while our results speak to the efficacy of the three studied attention mechanisms, there are numerous factors that we did not explore systematically in this study. For instance, it is well established that the degree of attentional modulation increases throughout the visual hierarchy, and is first observed in higher and later on in lower visual areas (Buffalo, Fries, Landman, Liang, & Desimone, 2010; Mehta, Ulbert, & Schroeder, 2000). Future studies using our modelling framework could study how these effects may dynamically come about. In particular, one could design different scenarios that independently vary the strength of attentional modulation, its timing and the targeted network layers to understand which combination may best match empirical results, and provide new insight into how attentional biases may be propagated through the visual hierarchy.

In this study, we examined how three main attention mechanisms can shape a complex and noisy process such as object recognition in natural scenes. Leveraging sDCNNs, we were able to inspect the computational consequences of different proposals of how selective attention may be implemented in the brain. Across a variety of measures, we observed a computational advantage of gain-based and in particular connection gain-based mechanisms, in contrast to precision. Our results highlight that sDCNNs provide a suitable modelling framework for connecting empirical observations from performance to neural processing and illustrate how they can be used to differentiate between theories.

## Acknowledgments

The authors would like to thank Leon Reteig for his advice on data visualization as well as several anonymous peer reviewers for their constructive feedback that helped to improve a past version of this manuscript.

## Notes

**Conflict of interest:** The authors declare no competing financial interests.

https://osf.io/6tpz8/

### Competing Interest Statement

The authors have declared no competing interest.

## References

Abadi, M., Agarwal, A., Barham, P., Brevdo, E., Chen, Z., Citro, C., … Zheng, X. (2016). TensorFlow: Large-Scale Machine Learning on Heterogeneous Distributed Systems. Retrieved from http://arxiv.org/abs/1603.04467

Allen, C., & Stevens, C. F. (1994). An evaluation of causes for unreliability of synaptic transmission. Proceedings of the National Academy of Sciences of the United States of America, 91(22), 10380–10383. doi:10.1073/pnas.91.22.10380

Anton-Erxleben, K., & Carrasco, M. (2013). Attentional enhancement of spatial resolution: linking behavioural and neurophysiological evidence. Nature Reviews. Neuroscience, 14(3), 188–200. doi:10.1038/nrn3443

Ayaz, A., & Chance, F. S. (2009). Gain modulation of neuronal responses by subtractive and divisive mechanisms of inhibition. Journal of Neurophysiology, 101(2), 958–968. doi:10.1152/jn.90547.2008

Bellec, G., Salaj, D., Subramoney, A., Legenstein, R., & Maass, W. (2018). Long short-term memory and Learning-to-learn in networks of spiking neurons. In S. Bengio, H. Wallach, H. Larochelle, K. Grauman, N. Cesa-Bianchi, & R. Garnett (Eds.), Advances in Neural Information Processing Systems 31 (pp. 787–797). Retrieved from http://papers.nips.cc/paper/7359-long-short-term-memory-and-learning-to-learn-in-networks-of-spiking-neurons.pdf

Benjamini, Y., & Hochberg, Y. (1995). Controlling the false discovery rate: a practical and powerful approach to multiple testing. Journal of the Royal Statistical Society. Retrieved from https://rss.onlinelibrary.wiley.com/doi/abs/10.1111/j.2517-6161.1995.tb02031.x

Beuth, F., & Hamker, F. H. (2015). A mechanistic cortical microcircuit of attention for amplification, normalization and suppression. Vision Research, 116(Pt B), 241–257. doi:10.1016/j.visres.2015.04.004

Bohte, S. (2012). Efficient Spike-Coding with Multiplicative Adaptation in a Spike Response Model. Advances in Neural Information Processing Systems25, 1844–1852. Retrieved from http://papers.nips.cc/paper/4698-efficient-spike-coding-with-multiplicative-adaptation-in-a-spike-response-model.pdf

Briggs, F., Mangun, G. R., & Usrey, W. M. (2013). Attention enhances synaptic efficacy and the signal-to-noise ratio in neural circuits. Nature, 499(7459), 476–480. doi:10.1038/nature12276

Buffalo, E. A., Fries, P., Landman, R., Liang, H., & Desimone, R. (2010). A backward progression of attentional effects in the ventral stream. Proceedings of the National Academy of Sciences of the United States of America, 107(1), 361–365. doi:10.1073/pnas.0907658106

Caesar, H., Uijlings, J., & Ferrari, V. (2016). COCO-Stuff: Thing and Stuff Classes in Context. Retrieved from http://arxiv.org/abs/1612.03716

Carrasco, M. (2011). Visual attention: the past 25 years. Vision Research, 51(13), 1484–1525. doi:10.1016/j.visres.2011.04.012

Chollet, F. (2015). keras. Retrieved from https://scholar.google.ca/scholar?cluster=17868569268188187229,14781281269997523089,11592651756311359484,12265559332197884258,14709450167780983337,17953590820456357796,6655887363479483357,5629189521449088544,11400611384887083769,5003160727454653660,10701427021387920284,694198723267881416&hl=en&as_sdt=0,5&sciodt=0,5

Cohen, M. R., & Maunsell, J. H. R. (2009). Attention improves performance primarily by reducing interneuronal correlations. Nature Neuroscience, 12(12), 1594–1600. doi:10.1038/nn.2439

Dayan, P., & Abbott, L. F. (2001). Theoretical neuroscience: computational and mathematical modeling of neural systems. Retrieved from https://pure.mpg.de/pubman/faces/ViewItemOverviewPage.jsp?itemId=item_3006127

Dayan, P., Kakade, S., & Montague, P. R. (2000). Learning and selective attention. Nature Neuroscience, 3 Suppl, 1218–1223. doi:10.1038/81504

Eickenberg, M., Gramfort, A., Varoquaux, G., & Thirion, B. (2017). Seeing it all: Convolutional network layers map the function of the human visual system. NeuroImage, 152, 184–194. doi:10.1016/j.neuroimage.2016.10.001

Feldman, H., & Friston, K. J. (2010). Attention, uncertainty, and free-energy. Frontiers in Human Neuroscience, 4, 215. doi:10.3389/fnhum.2010.00215

Ferguson, K. A., & Cardin, J. A. (2020). Mechanisms underlying gain modulation in the cortex. Nature Reviews. Neuroscience. doi:10.1038/s41583-019-0253-y

Galashan, F. O., Saßen, H. C., Kreiter, A. K., & Wegener, D. (2013). Monkey area MT latencies to speed changes depend on attention and correlate with behavioral reaction times. Neuron, 78(4), 740–750. doi:10.1016/j.neuron.2013.03.014

Güçlü, U., & van Gerven, M. A. J. (2015). Deep Neural Networks Reveal a Gradient in the Complexity of Neural Representations across the Ventral Stream. The Journal of Neuroscience: The Official Journal of the Society for Neuroscience, 35(27), 10005–10014. doi:10.1523/JNEUROSCI.5023-14.2015

Harris, C. R., Millman, K. J., van der Walt, S. J., Gommers, R., Virtanen, P., Cournapeau, D., … Oliphant, T. E. (2020). Array programming with NumPy. Nature, 585(7825), 357–362. doi:10.1038/s41586-020-2649-2

He, K., Zhang, X., Ren, S., & Sun, J. (2015). Deep Residual Learning for Image Recognition. Retrieved from http://arxiv.org/abs/1512.03385

Hembrook-Short, J. R., Mock, V. L., Usrey, W. M., & Briggs, F. (2018). Attention enhances the efficacy of communication in V1 local circuits. The Journal of Neuroscience: The Official Journal of the Society for Neuroscience. doi:10.1523/JNEUROSCI.2164-18.2018

Hunter, J. D. (2007). Matplotlib: A 2D Graphics Environment. Computing in Science Engineering, 9(3), 90–95. doi:10.1109/MCSE.2007.55

Itthipuripat, S., Cha, K., Byers, A., & Serences, J. T. (2017). Two different mechanisms support selective attention at different phases of training. PLoS Biology, 15(6), e2001724. doi:10.1371/journal.pbio.2001724

Itthipuripat, S., Ester, E. F., Deering, S., & Serences, J. T. (2014). Sensory gain outperforms efficient readout mechanisms in predicting attention-related improvements in behavior. The Journal of Neuroscience: The Official Journal of the Society for Neuroscience, 34(40), 13384–13398. doi:10.1523/JNEUROSCI.2277-14.2014

Jehee, J. F. M., Brady, D. K., & Tong, F. (2011). Attention improves encoding of task-relevant features in the human visual cortex. The Journal of Neuroscience: The Official Journal of the Society for Neuroscience, 31(22), 8210–8219. doi:10.1523/JNEUROSCI.6153-09.2011

Kar, K., Kubilius, J., Schmidt, K., Issa, E. B., & DiCarlo, J. J. (2019). Evidence that recurrent circuits are critical to the ventral stream’s execution of core object recognition behavior. Nature Neuroscience. Retrieved from https://www.nature.com/articles/s41593-019-0392-5

Khaligh-Razavi, S. M., & Kriegeskorte, N. (2014). Deep Supervised, but Not Unsupervised, Models May Explain IT Cortical Representation. PLoS Computational Biology, 10(11), e1003915. doi:10.1371/journal.pcbi.1003915

Kietzmann, T. C., McClure, P., & Kriegeskorte, N. (Eds.). (2019). Deep Neural Networks in Computational Neuroscience. In Oxford Research Encyclopedia of Neuroscience. doi:10.1093/acrefore/9780190264086.013.46

Kietzmann, T. C., Spoerer, C. J., Sörensen, L. K. A., Cichy, R. M., Hauk, O., & Kriegeskorte, N. (2019). Recurrence is required to capture the representational dynamics of the human visual system. Proceedings of the National Academy of Sciences of the United States of America, 116(43), 21854–21863. doi:10.1073/pnas.1905544116

Kriegeskorte, N., Mur, M., & Bandettini, P. A. (2008). Representational similarity analysis – connecting the branches of systems neuroscience. Frontiers in Systems Neuroscience, 2, 4. doi:10.3389/neuro.06.004.2008

Kubilius, J., Bracci, S., & Op de Beeck, H. P. (2016). Deep neural networks as a computational model for human shape sensitivity. PLoS Computational Biology, 12(4), e1004896. Retrieved from https://journals.plos.org/ploscompbiol/article?id=10.1371/journal.pcbi.1004896

Kubilius, J., Schrimpf, M., Nayebi, A., Bear, D., Yamins, D. L. K., & DiCarlo, J. J. (2018). CORnet: Modeling the Neural Mechanisms of Core Object Recognition (p. 408385). doi:10.1101/408385

Kümmerer, M., Wallis, T. S. A., & Bethge, M. (2016). DeepGaze II: Reading fixations from deep features trained on object recognition. Retrieved from http://arxiv.org/abs/1610.01563

Lazar, A. A., & Toth, L. T. (2003, April). Time encoding and perfect recovery of bandlimited signals. 2003 IEEE International Conference on Acoustics, Speech, and Signal Processing, 2003. Proceedings. (ICASSP ‘03)., 6, VI–709. doi:10.1109/ICASSP.2003.1201780

Lee, J., Williford, T., & Maunsell, J. H. R. (2007). Spatial attention and the latency of neuronal responses in macaque area V4. The Journal of Neuroscience: The Official Journal of the Society for Neuroscience, 27(36), 9632–9637. doi:10.1523/JNEUROSCI.2734-07.2007

Lin, T.-Y., Maire, M., Belongie, S., Bourdev, L., Girshick, R., Hays, J., … Dollár, P. (2014). Microsoft COCO: Common Objects in Context. Retrieved from http://arxiv.org/abs/1405.0312

Lindsay, G. W. (2020). Attention in Psychology, Neuroscience, and Machine Learning. Frontiers in Computational Neuroscience, 14, 29. doi:10.3389/fncom.2020.00029

Lindsay, G. W., & Miller, K. D. (2018). How biological attention mechanisms improve task performance in a large-scale visual system model. ELife, 7. doi:10.7554/eLife.38105

Lindsay, G. W., Rubin, D. B., & Miller, K. D. (2019). A simple circuit model of visual cortex explains neural and behavioral aspects of attention. Retrieved from https://www.biorxiv.org/content/biorxiv/early/2019/12/13/2019.12.13.875534.full.pdf

Luo, X., Roads, B. D., & Love, B. C. (2020). The Costs and Benefits of Goal-Directed Attention in Deep Convolutional Neural Networks. Retrieved from http://arxiv.org/abs/2002.02342

Ma, W. J., Beck, J. M., Latham, P. E., & Pouget, A. (2006). Bayesian inference with probabilistic population codes. Nature Neuroscience, 9(11), 1432–1438. doi:10.1038/nn1790

Martinez-Trujillo, J. C., & Treue, S. (2004). Feature-based attention increases the selectivity of population responses in primate visual cortex. Current Biology: CB, 14(9), 744–751. doi:10.1016/j.cub.2004.04.028

Martínez-Trujillo, J., & Treue, S. (2002). Attentional modulation strength in cortical area MT depends on stimulus contrast. Neuron, 35(2), 365–370. Retrieved from https://www.ncbi.nlm.nih.gov/pubmed/12160753

Maunsell, J. H. R. (2015). Neuronal Mechanisms of Visual Attention. Annual Review of Vision Science, 1, 373–391. doi:10.1146/annurev-vision-082114-035431

McAdams, C. J., & Maunsell, J. H. (1999). Effects of attention on orientation-tuning functions of single neurons in macaque cortical area V4. The Journal of Neuroscience: The Official Journal of the Society for Neuroscience, 19(1), 431–441. Retrieved from https://www.ncbi.nlm.nih.gov/pubmed/9870971

McKee, J. L., Riesenhuber, M., Miller, E. K., & Freedman, D. J. (2014). Task dependence of visual and category representations in prefrontal and inferior temporal cortices. The Journal of Neuroscience: The Official Journal of the Society for Neuroscience, 34(48), 16065–16075. doi:10.1523/JNEUROSCI.1660-14.2014

McKinney, W., & Others. (2010). Data structures for statistical computing in python. Proceedings of the 9th Python in Science Conference, 445, 51–56. Retrieved from http://conference.scipy.org/proceedings/scipy2010/pdfs/mckinney.pdf

Mehta, A. D., Ulbert, I., & Schroeder, C. E. (2000). Intermodal selective attention in monkeys. I: distribution and timing of effects across visual areas. Cerebral Cortex, 10(4), 343–358. doi:10.1093/cercor/10.4.343

Mitchell, J. F., Sundberg, K. A., & Reynolds, J. H. R. (2007). Differential attention-dependent response modulation across cell classes in macaque visual area V4. Neuron, 55(1), 131–141. doi:10.1016/j.neuron.2007.06.018

Mitchell, J. F., Sundberg, K. A., & Reynolds, J. H. R. (2009). Spatial attention decorrelates intrinsic activity fluctuations in macaque area V4. Neuron, 63(6), 879–888. doi:10.1016/j.neuron.2009.09.013

Nair, V., & Hinton, G. E. (2010). Rectified Linear Units Improve Restricted Boltzmann Machines. Retrieved from https://openreview.net/pdf?id=rkb15iZdZB

Naka, K.-I., & Rushton, W. A. H. (1966). S-potentials from luminosity units in the retina of fish (Cyprinidae). The Journal of Physiology, 185(3), 587–599. Retrieved from https://physoc.onlinelibrary.wiley.com/doi/abs/10.1113/jphysiol.1966.sp008003

Parr, T., & Friston, K. J. (2017). Working memory, attention, and salience in active inference. Scientific Reports, 7(1), 14678. doi:10.1038/s41598-017-15249-0

Pedregosa, F. (2011). Scikit-learn: Machine Learning in Python. Journal of Machine Learning Research: JMLR, 12, 2825–2830. Retrieved from https://jmlr.csail.mit.edu/papers/volume12/pedregosa11a/pedregosa11a.pdf

Posner, M. I. (1980). Orienting of attention. The Quarterly Journal of Experimental Psychology, 32(1), 3–25. Retrieved from https://www.ncbi.nlm.nih.gov/pubmed/7367577

Reynolds, J. H. R., & Heeger, D. J. (2009). The normalization model of attention. Neuron, 61(2), 168–185. doi:10.1016/j.neuron.2009.01.002

Reynolds, J. H. R., Pasternak, T., & Desimone, R. (2000). Attention increases sensitivity of V4 neurons. Neuron, 26(3), 703–714. Retrieved from https://www.ncbi.nlm.nih.gov/pubmed/10896165

Richards, B. A., Lillicrap, T. P., Beaudoin, P., Bengio, Y., Bogacz, R., Christensen, A., … Kording, K. P. (2019). A deep learning framework for neuroscience. Nature Neuroscience, 22(11), 1761–1770. doi:10.1038/s41593-019-0520-2

Rothenstein, A. L., & Tsotsos, J. K. (2014). Attentional modulation and selection--an integrated approach. PloS One, 9(6), e99681. doi:10.1371/journal.pone.0099681

Rueckauer, B., Lungu, I.-A., Hu, Y., & Pfeiffer, M. (2016). Theory and Tools for the Conversion of Analog to Spiking Convolutional Neural Networks. Retrieved from http://arxiv.org/abs/1612.04052

Russakovsky, O., Deng, J., Su, H., Krause, J., Satheesh, S., Ma, S., … Fei-Fei, L. (2015). ImageNet Large Scale Visual Recognition Challenge. International Journal of Computer Vision, 115(3), 211–252. doi:10.1007/s11263-015-0816-y

Scholte, H. S. (2018). Fantastic DNimals and where to find them. NeuroImage, 180(Pt A), 112–113. doi:10.1016/j.neuroimage.2017.12.077

Scholte, H. S., Ghebreab, S., Waldorp, L., Smeulders, A. W. M., & Lamme, V. A. F. (2009). Brain responses strongly correlate with Weibull image statistics when processing natural images. Journal of Vision, 9(4), 29.1-15. doi:10.1167/9.4.29

Schrimpf, M., Kubilius, J., Lee, M. J., Ratan Murty, N. A., Ajemian, R., & DiCarlo, J. J. (2020). Integrative Benchmarking to Advance Neurally Mechanistic Models of Human Intelligence. Neuron. doi:10.1016/j.neuron.2020.07.040

Seijdel, N., Tsakmakidis, N., de Haan, E. H. F., Bohte, S. M., & Scholte, H. S. (2020). Depth in convolutional neural networks solves scene segmentation. PLoS Computational Biology, 16(7), e1008022. doi:10.1371/journal.pcbi.1008022

Spoerer, C. J., Kietzmann, T. C., Mehrer, J., Charest, I., & Kriegeskorte, N. (2020). Recurrent neural networks can explain flexible trading of speed and accuracy in biological vision. PLoS Computational Biology, 16(10), e1008215. doi:10.1371/journal.pcbi.1008215

Sundberg, K. A., Mitchell, J. F., Gawne, T. J., & Reynolds, J. H. R. (2012). Attention influences single unit and local field potential response latencies in visual cortical area V4. The Journal of Neuroscience: The Official Journal of the Society for Neuroscience, 32(45), 16040–16050. doi:10.1523/JNEUROSCI.0489-12.2012

Treue, S., & Martínez Trujillo, J. C. (1999). Feature-based attention influences motion processing gain in macaque visual cortex. Nature, 399(6736), 575–579. doi:10.1038/21176

Vallat, R. (2018). Pingouin: statistics in Python. Journal of Open Source Software, 3(31), 1026. doi:10.21105/joss.01026

VanRullen, R. (2017). Perception Science in the Age of Deep Neural Networks. Frontiers in Psychology, 8, 142. doi:10.3389/fpsyg.2017.00142

Virtanen, P., Gommers, R., Oliphant, T. E., Haberland, M., Reddy, T., Cournapeau, D., … SciPy 1.0 Contributors. (2020). SciPy 1.0: fundamental algorithms for scientific computing in Python. Nature Methods, 17(3), 261–272. doi:10.1038/s41592-019-0686-2

Waskom, M., Botvinnik, O., Gelbart, M., Ostblom, J., Hobson, P., Lukauskas, S., … Brunner, T. (2020). mwaskom/seaborn: v0.11.0 (Sepetmber 2020). doi:10.5281/zenodo.4019146

Wyart, V., Nobre, A. C., & Summerfield, C. (2012). Dissociable prior influences of signal probability and relevance on visual contrast sensitivity. Proceedings of the National Academy of Sciences of the United States of America, 109(9), 3593–3598. doi:10.1073/pnas.1120118109

Yamins, D. L. K., & DiCarlo, J. J. (2016). Using goal-driven deep learning models to understand sensory cortex. Nature Neuroscience, 19(3), 356–365. doi:10.1038/nn.4244

Yamins, D. L. K., Hong, H., Cadieu, C. F., Solomon, E. A., Seibert, D., & DiCarlo, J. J. (2014). Performance-optimized hierarchical models predict neural responses in higher visual cortex. Proceedings of the National Academy of Sciences of the United States of America, 111(23), 8619–8624. doi:10.1073/pnas.1403112111

Yoon, Y. C. (2017). LIF and Simplified SRM Neurons Encode Signals Into Spikes via a Form of Asynchronous Pulse Sigma-Delta Modulation. IEEE Transactions on Neural Networks and Learning Systems, 28(5), 1192–1205. doi:10.1109/TNNLS.2016.2526029

Yu, A. J., & Dayan, P. (2005). Uncertainty, neuromodulation, and attention. Neuron, 46(4), 681–692. doi:10.1016/j.neuron.2005.04.026

Zambrano, D., Nusselder, R., Scholte, H. S., & Bohté, S. M. (2018). Sparse Computation in Adaptive Spiking Neural Networks. Frontiers in Neuroscience, 12, 987. doi:10.3389/fnins.2018.00987

Zhang, Y., Meyers, E. M., Bichot, N. P., Serre, T., Poggio, T. A., & Desimone, R. (2011). Object decoding with attention in inferior temporal cortex. Proceedings of the National Academy of Sciences of the United States of America, 108(21), 8850–8855. doi:10.1073/pnas.1100999108

